# High-precision cell-type mapping and annotation of single-cell spatial transcriptomics with STAMapper

**DOI:** 10.1101/2025.01.08.631859

**Authors:** Qunlun Shen, Kangning Dong, Shuqin Zhang, Shihua Zhang

**Affiliations:** School of Mathematical Sciences, Fudan University, Shanghai, 200433, China; School of Mathematics, Renmin University of China, Beijing 100872, China; NCMIS, CEMS, RCSDS, Academy of Mathematics and Systems Science, Chinese Academy of Sciences, Beijing 100190, China; Key Laboratory of Mathematics for Nonlinear Science, Fudan University, Ministry of Education, Shanghai, 200433, China; Shanghai Key Laboratory for Contemporary Applied Mathematics, Fudan University, Shanghai, 200433, China; School of Mathematical Sciences, University of Chinese Academy of Sciences, Beijing 100049, China; Key Laboratory of Systems Health Science of Zhejiang Province, School of Life Science, Hangzhou Institute for Advanced Study, University of Chinese Academy of Sciences, Chinese Academy of Sciences, Hangzhou 310024, China

## Abstract

Recent advances in single-cell spatial transcriptomics (scST) have enabled the analysis of gene transcription levels in individual cells while preserving their spatial positions. Cell-type mapping and annotation are crucial in understanding the complex interactions between cells and their microenvironments within a spatial context. To this end, we develop a heterogeneous graph neural network, STAMapper, to transfer the cell-type labels from single-cell RNA-seq data to scST data. STAMapper captures both the expression similarity among cells and the expression relationships between cells and genes and adopts a graph attention classifier to conduct semi-supervised learning for more accurate cell-type prediction. We collected 81 scST datasets consisting of 344 slices and 16 paired scRNA-seq datasets from eight technologies and five tissues to validate the efficiency of STAMapper. STAMapper achieved the best performance on 75 out of 81 datasets compared to competing methods in accuracy. STAMapper demonstrated enhanced performance over manual annotations, particularly at the boundaries of cell clusters, enabled the unknown cell-type detection in scST data, and exhibited precise cell subtype annotations. Additionally, STAMapper provided biologically meaningful gene embeddings, facilitating the identification of shared or unique gene modules across datasets.

## Introduction

Single-cell RNA sequencing (scRNA-seq) technologies allow us to study whole-transcriptional changes at the single-cell level to shape the diversity of cell types and their dynamic changes^1^. However, the spatial position information of single cells tends to be lost due to the dissociation during the sequencing process, which prevents us from understanding the relationship between gene expression and the tissue architecture and hinders us from deciphering the complex interactions between cells and their microenvironments under spatial context^2^. More recently, the emergence of single-cell spatial transcriptomics (scST) technologies such as MERFSIH^3, 4^, seqFISH^5^, seqFISH+^6^, osmFISH^7^, STARmap^8^, STARmap PLUS^9^, NanoString^10^, and Slide-tags^11^, enables the profiling of gene expression with their spatial context at single-cell resolution.

The essential problem in scRNA-seq and scST data analysis is cell-type annotation (or cell typing)^12, 13^. The standard workflow^14^ for cell-type annotation in scRNA-seq data is normalization, gene selection (usually top 2000 highly variable genes), dimensionality reduction, clustering, and assigning a cell type to each cluster according to the expression of known marker genes. When dealing with the scST data, the above workflow may fail since the sequencing quality of scST technologies is far lower than that of the mature scRNA-seq technologies. Specifically, these spatially resolved technologies typically focus on a pre-defined set of marker genes or genes relevant to biological processes (usually far fewer than 2000, **Table S1**). In the case of Slide-tags, a whole-transcriptome single nucleus spatial technology, approximately 75% of nuclei are lost during sequencing^11^. These factors may lead to clustering instability and blurred cluster boundaries, resulting in inaccurate cell-type annotation. Moreover, as some of the markers for rare cell types may be absent in ST data, the annotation for the related cells could be challenged or overlooked. Therefore, accurate annotation of single-cell ST data remains demanding and intricate.

With more scRNA-seq data available, reference-based scST annotation methods have been proposed to transfer cell-type labels to query datasets by leveraging the well-annotated reference dataset^15–17^. For example, scANVI employs a variational autoencoder architecture to learn a latent space of cellular states for both the scRNA-seq and scST data and utilizes the mean of the variational distribution associated with each cell to perform annotation^15^. RCTD utilizes a regression framework to model cell-type profiles in reference and account for platform effects, facilitating cell-type identification in spatial data^16^. Tangram maps scRNA-seq profiles onto ST data by maximizing the cosine similarity of the predicted and the observed expression matrix^17^. These methods can predict cell-type labels in datasets generated by MERFISH (scANVI)^18^, Slide-seq (RCTD)^16^, and STARmap (Tangram)^17^. However, these existing methods may fail to reveal fuzzy boundaries in scST annotations, and due to the lack of incorporating gene modeling, they could not identify gene modules either shared by scRNA-seq and scST data or unique to each of them. Furthermore, to the best of our knowledge, there has not yet been a substantial number of prepared real datasets to evaluate the accuracy and robustness of the cell-type annotation methods across different ST technologies and tissue origins.

To this end, we develop STAMapper to accurately annotate cells from scST data by a heterogeneous graph neural network^19^ with a graph attention classifier. Also, we collected 81 paired scRNA-seq and scST datasets with manual annotations and manually aligned them to evaluate the annotation accuracy. Extensive tests and comparisons with existing methods demonstrated the superiority of STAMapper in various biological applications, i.e., cell-type mapping of scST data, reannotation of blurred cell types, unknown cell-type detection, subtype annotation, and gene module extraction. Additionally, the collected data can serve as a benchmark for scST annotation, and the annotation results of STAMapper can serve as a reference.

## Results

### Overview of STAMapper

STAMapper takes a well-annotated scRNA-seq dataset and a scST dataset as input, where the two data matrices are normalized. STAMapper first constructs a heterogeneous graph, where the cells and genes are modeled as two distinct types of nodes and connected with edges based on whether the genes are expressed in the cells. Two cells from each dataset are connected if they exhibit similar gene expression patterns. Each node is connected to itself to indicate it utilizes the information from the previous step when updating its embedding (**Fig. 1**, **Methods**).

**Figure 1.**
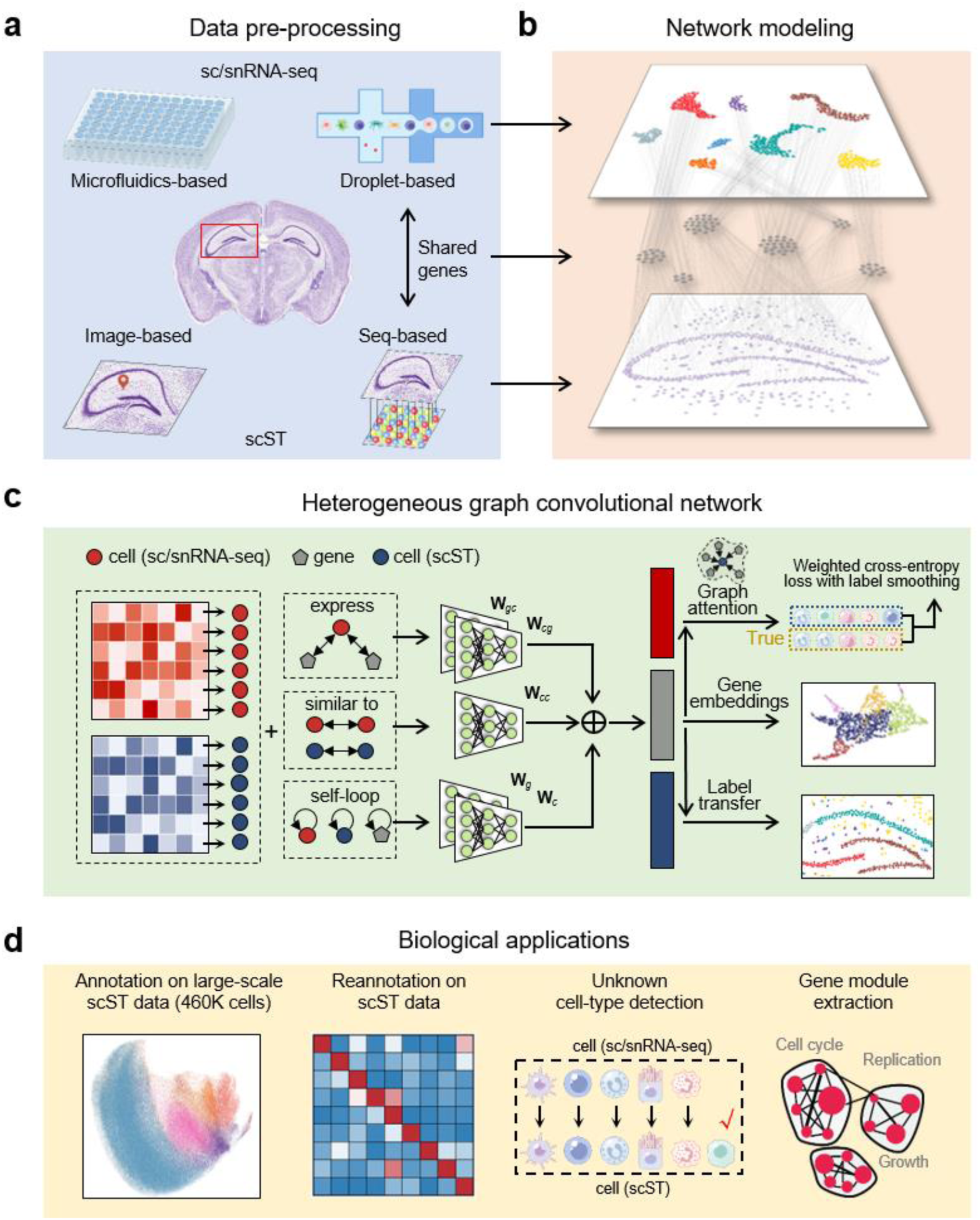
Illustration of STAMapper and its applications. **a.** STAMapper can annotate scST data obtained from mainstream technologies such as image-based and seq-based by leveraging well-annotated sc/snRNA-seq data sequenced from microfluidics-based or droplet-based technologies. **b.** STAMapper models genes and cells as two types of heterogeneous nodes and connects sc/scRNA-seq and scST data by their expression on the shared genes. **c.** STAMapper takes the expression and the heterogeneous relationships of nodes as input. STAMapper then learns embeddings for cells and genes based on the information propagation mechanism on the heterogeneous graph network to fit cell labels from scRNA-seq data by using a graph attention classifier, ultimately utilizing the learned weights of information propagation on the graph to transfer cell labels on spatial data. **d.** The output of STAMapper can be applied for annotation on large-scale scST data, reannotation on scST data, unknown cell-types detection, and gene module extraction.

For each cell node, the initial input is the corresponding normalized gene expression vector. The gene nodes obtain their initial embedding by aggregating the input from the connected cell nodes (**Methods**). STAMapper updates the latent embedding of each cell or gene node based on the message-passing mechanism with information from their neighbors. It utilizes the embedding of gene nodes as the input of a graph attention classifier to estimate the probability of the cell-type identity, wherein each cell assigns varying attention weights to its connected genes. STAMapper uses a modified cross-entropy loss^20^ (**Methods**) to quantify the discrepancy between the predicted and original cell-type labels for cells in the scRNA-seq dataset. Finally, through backpropagation, STAMapper updates the weights of parameters for different edges until the model converges. STAMapper determines gene modules based on the learned embeddings of gene nodes with the Leiden clustering algorithm^21^ and applies the outputs of the graph attention classifier to assign cell-type labels to cells in the scST dataset.

### STAMapper enables accurate cell-type mapping for scST data

We collected 81 single-cell ST datasets comprised of 344 slices and 16 paired scRNA-seq datasets from identical tissues. These scST datasets originate from eight single-cell ST technologies, i.e., MERFISH^3^, NanoString^10^, STARmap^8^, STARmap Plus^9^, Slide-tags^11^, osmFISH^7^, seqFISH^5^, seqFISH+^6^, and five different tissues, i.e., brain, embryo, retina, kidney, liver (**Fig. 2a**). All datasets come with manual annotations provided by the authors and the cell-type labels in paired scRNA-seq and spatial datasets aligned manually.

**Figure 2.**
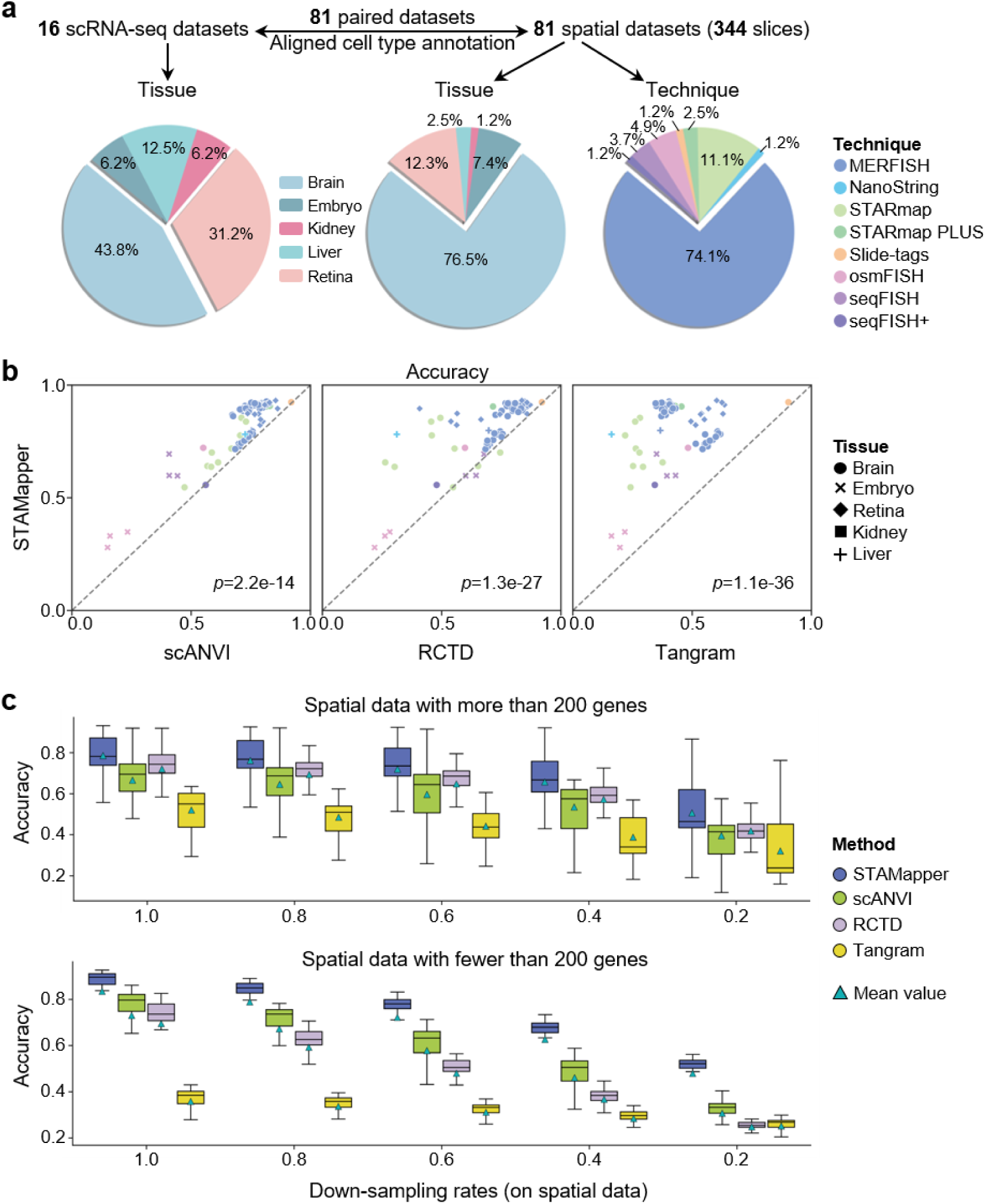
Benchmarking cell annotation performance of STAMapper. **a.** Overview of all datasets used for evaluating the performance of STAMapper. We collected 81 single-cell spatial transcriptomics datasets comprising a total of 344 slices, where each dataset is matched with corresponding single-cell transcriptomics data (or scRNA-seq data). **b.** Performance comparison of STAMapper and scANVI, RCTD, Tangram regarding cell annotation accuracy on 81 pairs of scRNA-seq and single-cell spatial transcriptomics datasets. *P*-values were calculated by paired t-test. **c.** Performance comparison of the classification accuracies of STAMapper and three other methods on different down-sampling rates (1.0, 0.8, 0.6, 0.4, 0.2) for read counts, where the down-sampling rate of 1.0 means the raw data. The upper panel depicts spatial transcriptomics datasets with more than 200 genes for sequencing (47 datasets), while the lower panel corresponds to fewer than 200 genes (34 datasets).

We quantitatively evaluated the cell-type annotation performance of STAMapper and the competing methods, i.e., scANVI^15^, RCTD^16^, and Tangram^17^, in terms of accuracy, macro F1 score, and weighted F1 score (**Methods**). STAMapper demonstrated significantly higher accuracy in annotating cells from scST datasets compared with scANVI (*p*=2.2e-14), RCTD (*p*=1.3e-27), and Tangram (*p*=1.3e-36) (**Fig. 2b**). It is noteworthy that, STAMapper achieved the best performance on 75 out of 81 datasets compared to competing methods in terms of accuracy. It also achieved the best overall performance with the macro F1 score and weighted F1 score for the imbalanced cell-type distributions (**Supplementary Fig. 1a, b**). scANVI ranked the second best across the three metrics. These results suggest that STAMapper exhibits the best annotation capability and proficiently identifies the rare cell types, which is crucial in cell-type annotation.

We evaluated the performance under poor sequencing quality with four different down-sampling rates on scST data. STAMapper consistently demonstrated the highest accuracy, macro F1 score, and weighted F1 score (**Fig. 1c**, **Supplementary Fig. 1c, d**). This trend is particularly distinct in scST datasets with fewer than 200 genes, where at a down-sampling rate of 0.2, STAMapper exhibited a much higher accuracy than the second-highest ranking method scANVI (median 51.6% VS 34.4%) (**Fig. 1c**). RCTD demonstrated superiority in the raw data (25 of 34 datasets) and comparable performance in the down-sampled data (**Fig. 1c**) for datasets containing more than 200 genes. scANVI tended to outperform RCTD on scST datasets with fewer than 200 genes in the raw data (41 of 47 datasets) and the down-sampled data regarding the accuracy and weighted F1 score. scANVI showed comparable performance with RCTD for datasets containing more than 200 genes, consistently outperforming RCTD on scST datasets with fewer than 200 genes in terms of macro F1 score (**Supplementary Fig. 1c, d**). Additionally, to mitigate the sensitivity of deep learning methods to parameters, we tested the accuracy of scANVI under various parameters (**Supplementary Fig. 1e**).

### STAMapper facilitates precise cell-type mapping within the retinal laminar structure

We applied STAMapper to ten MERFISH datasets derived from the mouse retina^22^, a highly organized tissue, on which we assess the accuracy of annotations from the perspective of cell spatial positions beyond checking the expression of markers. Here, we selected five scRNA-seq datasets of the mouse retina collected from postnatal 0 hours to 60 days (P60) as the reference data^23^ to test the robustness of STAMapper. As expected, STAMapper consistently outperformed the other three methods on every spatial dataset according to accuracy, macro F1 score, and weighted F1 score (**Fig. 3a**, **Supplementary Fig. 2a-c**). Additionally, STAMapper exhibited the lowest variance in accuracy and weighted F1 score, demonstrating its robustness to changes in the reference data. Due to the poor performance of Tangram, we did not include it in subsequent analyses.

**Figure 3.**
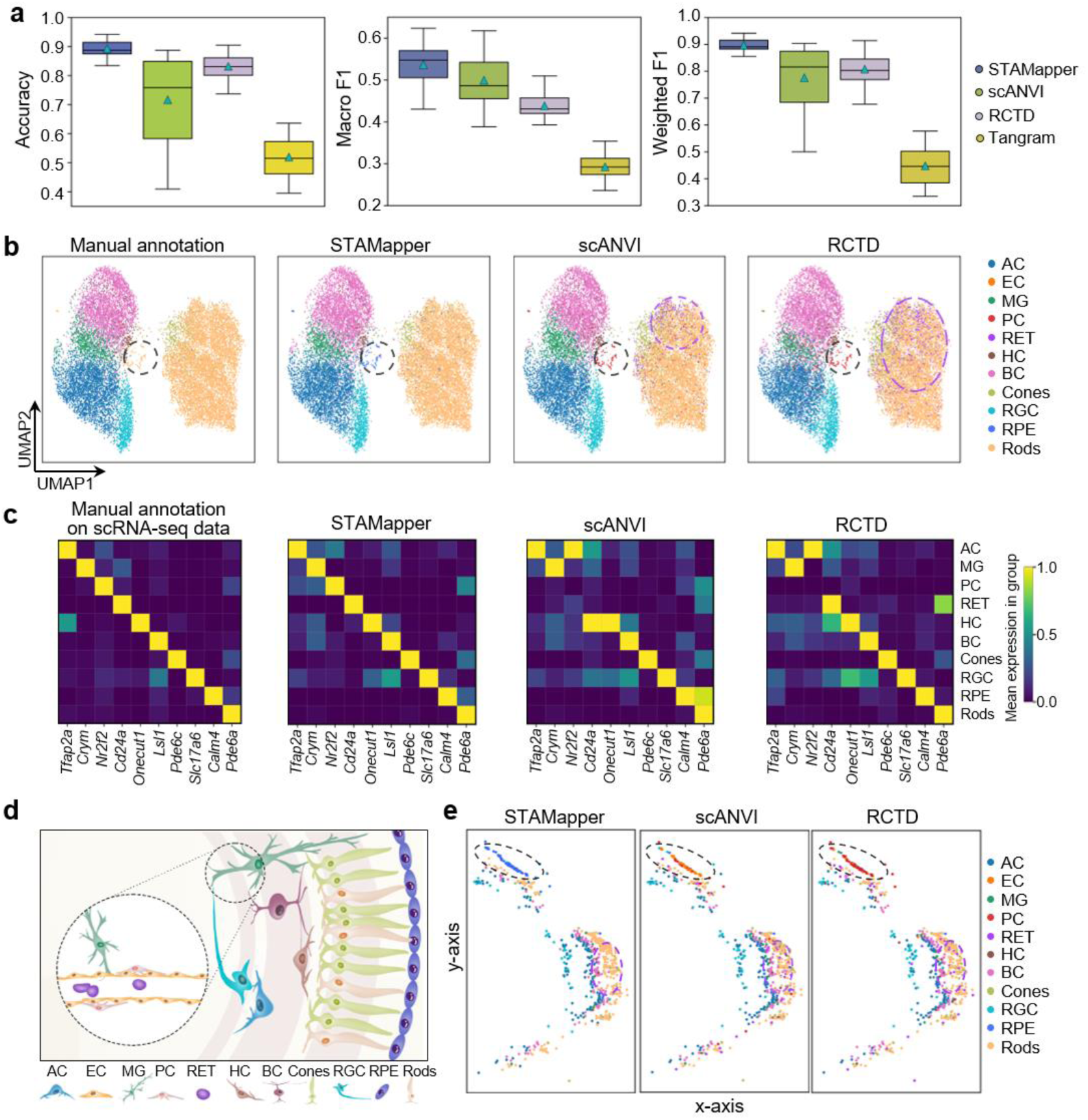
Application of STAMapper to MERFISH retina datasets. **a.** Performance comparison of STAMapper and scANVI, RCTD, Tangram, where each box represents the method’s performance on the 50 paired datasets (five scRNA-seq datasets and ten single-cell spatial transcriptomics datasets). **b.** UMAP plots of mouse retinal dataset (VZG105a_WT3) cells colored by the manual annotation and the prediction of STAMapper, scANVI, and RCTD using the mouse_LD_60 scRNA-seq dataset as the reference. AC, amacrine cells; EC, endothelial cells; MG, Müller Glia; PC, pericytes; RET, reticulocyte; HC, retinal horizontal cells; BC, bipolar cells; Cones, cone cells; RGC, retinal ganglion cells; RPE, retinal pigment epithelium; Rods, Rod cells. **c.** The heatmap of the marker expression for major cell types on the scRNA-seq dataset grouped by manual annotation and on the corresponding spatial transcriptomics dataset annotated by STAMapper, scANVI, and RCTD, respectively. **d.** A schematic illustration of the distribution of cell types within the retina. **e.** Spatial organization of a slice from spatial transcriptomics dataset corresponding to **(b)**, where cells are colored by the annotation by STAMapper, scANVI, and RCTD, respectively.

ScRNA-seq often allows for identifying some rare cell types since it can measure the expressions of most genes. However, some cell types identified in single-cell data, including endothelial cells (EC), pericytes (PC), retinal pigment epithelium (RPE), and reticulocyte (RET), do not appear in the manual annotations of spatial data, possibly due to the limited number of genes sequenced (**Table S1**) or inadequacies in clustering methods. In particular, the considered spatial data only contained 368 genes, where not all the markers^23^ of the cell types in scRNA-seq data were captured. In this case, STAMapper, scANVI, and RCTD were able to annotate some of these cell types. STAMapper annotated a group of rods (black dotted line in **Fig. 3b**), distant from the major cluster, as RPE but as PC by scANVI and RCTD. Additionally, scANVI and RCTD identified a cluster of RETs (purple dotted line in **Fig. 3b**) at the top of the rods cluster. To determine whether the annotated cell types indeed exist in the scST data, we first selected markers^23^ in original spatial data and differential expressed genes for the newly annotated cell types and further validated by the scRNA-seq data (**Fig. 3c**). The expression of these genes confirmed the accurate annotation of STAMapper. The PCs annotated by scANVI and RCTD did not express *Nr2f2*, and the RETs also demonstrated inconsistency in the expression of *Cd24a,* which were two genes showing highly differential expression in the corresponding cell types from scRNA-seq data (**Fig. 3c**). This indicated that there could be inaccuracies in their annotation.

We further examined their spatial locations to validate the annotation of newly discovered cell types. Here, scANVI and RCTD identified the cells in the outermost layer of the retina as ECs and PCs (black dashed ellipse in **Fig. 3e**), which does not align well with their real anatomical location^24, 25^ (the inner layer of retina) (**Fig. 3d**), while STAMapper annotated this group of cells as RPE cells, which expressed the corresponding markers in the appropriate anatomical positions^26^ (**Fig. 3c-e**). Different from STAMapper, scANVI and RCTD mistakenly annotated PCs among the rods (black dashed ellipse in Fig.3e), which should be located in the retina’s inner side, consistent with our observations on UMAP (**Fig. 3b**). In addition, scANVI and RCTD wrongly identified many RET cells between rods (purple dashed ellipse in **Fig. 3e**), which should be located at the most inner part of the retina (**Fig. 3d**). Similar situations can be observed across the other six slices (**Supplementary Fig. 2d-i**). Overall, STAMapper uncovered the cell types that clustering alone fails to identify in spatial retinal data by making full use of the detailed annotations from scRNA-seq data. More importantly, the annotation of STAMapper perfectly aligned with the retinal architecture, spanning from the outer retinal cells to the inner supporting cells.

### STAMapper corrects the blurred cell-type annotations caused by clustering-induced boundaries

We further applied STAMapper to the MERFISH hypothalamic data (ID=15) collected from mouse brain^27^ with the reference scRNA-seq data from the same study to test whether it can help correct the cell-type annotations at the boundaries detected by clustering algorithms. STAMapper achieved the highest annotation accuracy (86.3%) compared with scANVI (72.6%) and RCTD (67.3%). Specifically, scANVI mistakenly identified many macrophages mixed with inhibitory neurons and some fibroblasts at the boundary of OD mature cells. RCTD mistakenly annotated some ependymal cells mixed with inhibitory neurons and excitatory cells. In contrast, the annotation of STAMapper avoided the mixing of cell types between two non-transitional cells and showed smoothness at the boundaries of the clusters (**Fig. 4a**). Taking a close look at a slice of this sample, we discovered that STAMapper accurately recovered the “arrow-like” excitatory neurons in the middle of the slice, whereas RCTD mistakenly annotated them as a mixture of excitatory neurons and ependymal cells. STAMapper correctly identified the cellular microenvironment dominated by inhibitory neurons, and scANVI and RCTD failed (**Fig. 4b**). For the remaining three slices, STAMapper maintained the highest degree of consistency with manual annotations (**Supplementary Fig. 3a-c**). The Sankey plot of the entire sample demonstrated that scANVI and RCTD incorrectly predicted many inhibitory neurons to other cells like astrocytes, macrophages, and excitatory neurons. (**Fig. 4c**). scANVI predicted a considerable number of macrophages annotated as inhibitory neurons manually. However, these cells did express the inhibitory neuron marker *Gad1*, indicating the annotation errors by scANVI (**Supplementary Fig. 3d**).

**Figure 4.**
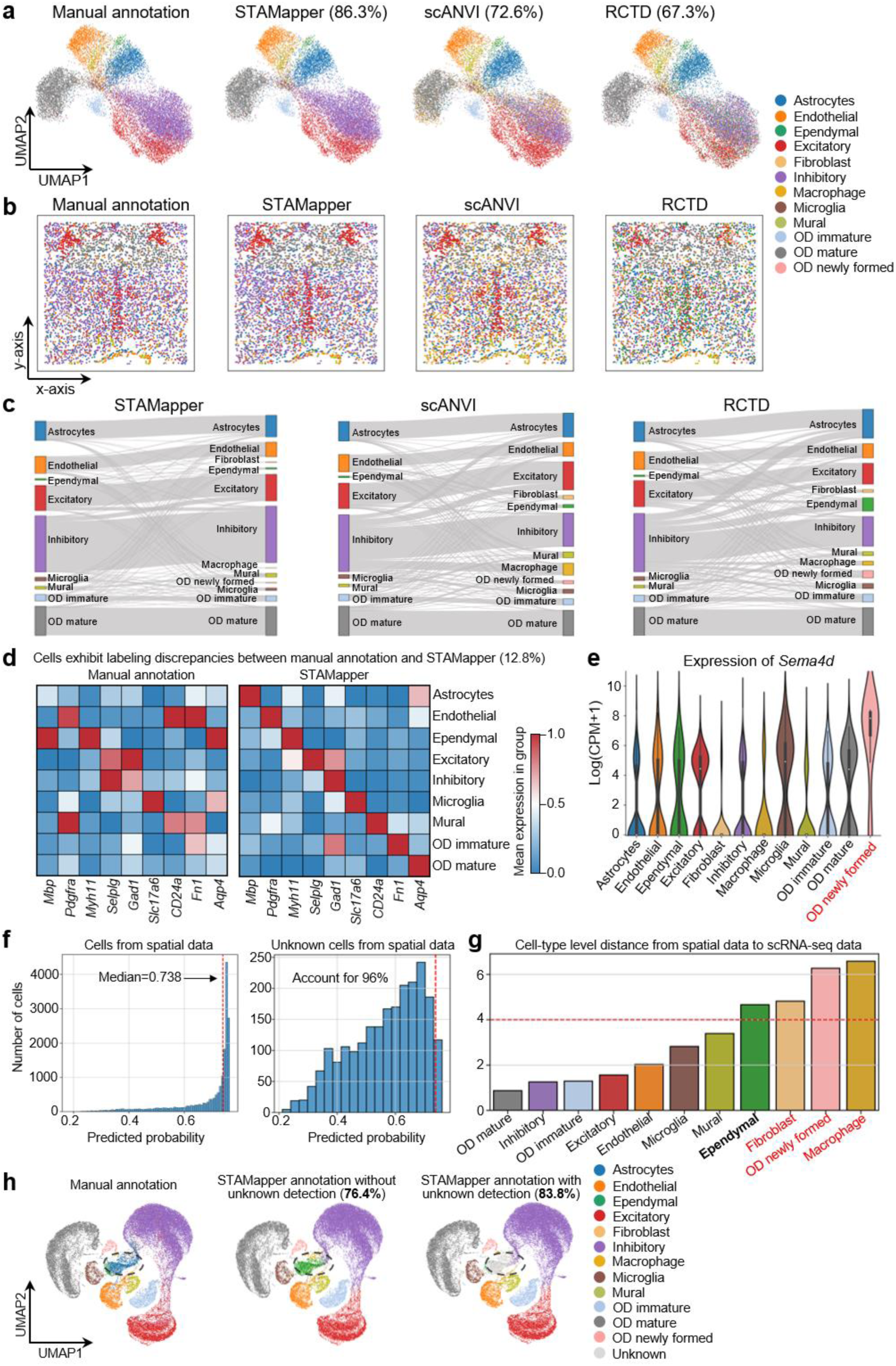
Application of STAMapper to MERFISH hypothalamic dataset. **a.** UMAP plots of mouse hypothalamic dataset colored by the manual annotation and the prediction of STAMapper, scANVI, and RCTD, respectively. **b.** Spatial organization of a slice from mouse hypothalamic dataset corresponding to **(a)**, cells are colored by the manual annotation and the prediction of STAMapper, scANVI, and RCTD, respectively. **c.** Sankey plot showing the accuracy of the cell-type annotation by STAMapper, scANVI, and RCTD, respectively. The left side of the Sankey plot represents manual annotations, while the right side shows the predicted results. The height of each linkage line reflects the number of cells. **d.** Heatmap plot of marker expression for major cell types presented in manual annotation with mismatched labels between manual annotation and STAMapper. **e.** Expression levels of *Sema4d* (a marker of OD Newly formed) across different cell types (annotated by STAMapper). **f.** The predicted probability of STAMapper for cells from spatial data (left panel) and unknown cells from spatial data (right panel), the red dash line indicates x=0.738 in both panels. **g.** Cell-type level distance from spatial data to single-cell data on cell embeddings learned by STAMapper. Bold indicates unknown cells were predicted as this specific cell type, and red denotes cell types present in single-cell data but not annotated in spatial data by manual annotation. **h.** UMAP plots of the co-embedding of scRNA-seq and spatial data learned by STAMapper, cells are colored by manual annotation, STAMapper prediction without unknown detection, and STAMapper prediction with unknown detection. The percentages in parentheses represent the predicted accuracy.

Despite STAMapper exhibiting the highest annotation accuracy, some cells showed inconsistencies with manual annotations. These cells tended to be located at the boundaries of cell clusters, e.g., around microglia (**Fig. 4a**). We examined the cell types already presented in the manual annotations. Some cell types didn’t express the corresponding markers from the original study^27^. Yet the annotation of STAMapper aligned precisely with the expression of marker genes (**Fig. 4d**). Also, STAMapper identified OD newly formed cells, which were not previously recognized in manual annotations, verified by highly expressing their marker gene *Sema4d*^28^ (**Fig. 4e**).

We further extended STAMapper to detect unknown cell types in spatial data. Here, we removed astrocytes from the scRNA-seq dataset and assumed to identify them as unknown cells in the spatial data. We defined the potentially unknown cells by the following two rules: (i) predicted probability less than the median value of all cells from spatial data, and (ii) the cell-type level distance from scST data to scRNA-seq data higher than a user-defined threshold (i.e., 4). (**Fig. 4f, g**, **Methods**). In this scenario, STAMapper without unknown cell detection primally achieved an accuracy of 76.4%, where it annotated astrocytes as ependymal reasonably, which is another type of glial cell. Leveraging the mechanism of unknown detection, STAMapper successfully annotated astrocytes as unknown cells and increased the accuracy to 83.8% (**Fig. 4h**). We also performed another test, where we removed OD immature cells from the scRNA-seq dataset, and STAMapper continued to annotate these cells as unknown correctly (**Supplementary Fig. 3e-g**). Therefore, STAMapper could rectify cell types at classification boundaries that are challenging to identify with clustering algorithms and detect unknown cell types in spatial data.

### STAMapper achieves precise annotations aiding in deciphering the tumor microenvironment

We applied STAMapper to the human hepatocellular carcinoma spatial data sequenced by NanoString technology^10^, and the scRNA-seq data originated from the same type of tumor lesions as a reference^29^. Compared to scANVI and RCTD, the annotation of STAMapper could reveal distinct boundaries between different cell types and achieve the highest consistency with the annotations provided by the authors (**Fig. 5a, b**). The expression of markers aligned closely with the corresponding cell types^30^, i.e., *PROX1* (Malignant), *CD163* (Macro), *NKG7* (NK), *CD3D* (T cell), *CD8A* (CD8 T), *IL7R* (CD4 T), *PECAM1* (Endothelial), *COL1A1* (Fibroblast), *JCHAIN* (Mature B) (**Fig.5a, c**). We selected two regions of interest (ROIs) in the tissue section (**Supplementary Fig. 4a**). ROI 1 was mainly malignant cells marked with highly expressed *PROX1*. scANVI annotated more DCs in this region, while RCTD annotated some cholangiocytes. However, the markers corresponding to these two cell types are barely expressed in this region (**Supplementary Fig. 4b, c**). ROI 2 appeared as a region mixed with immune cells and malignant cells, where numerous cells are distributed in a circular formation in the middle of this region (**Fig. 5d**). STAMapper annotated it as a microenvironment where macrophages enveloped malignant cells, T cells located in the outer side of this structure, and the exterior layer consisted of mature B cells. RCTD annotated the innermost layer with cholangiocytes surrounded by malignant cells, whereas scANVI annotated the innermost layer as mainly cholangiocytes with very few malignant cells. The expression of *PROX1* validated the presence of malignant cells in the innermost layer, and its expression is consistent with the annotations of STAMapper (**Fig. 5d, e**). Meanwhile, cells in this region barely expressed *KRT7*, a marker of cholangiocyte^31^, indicating that the annotations by RCTD and scANVI were incorrect (**Fig. 5e**). Interestingly, all three methods agreed that there exists a reticular structure of macrophages in this area (**Supplementary Fig. 4d**), and in fact, they surrounded the malignant cells, forming a unique microenvironment (**Fig. 5d**).

**Figure 5.**
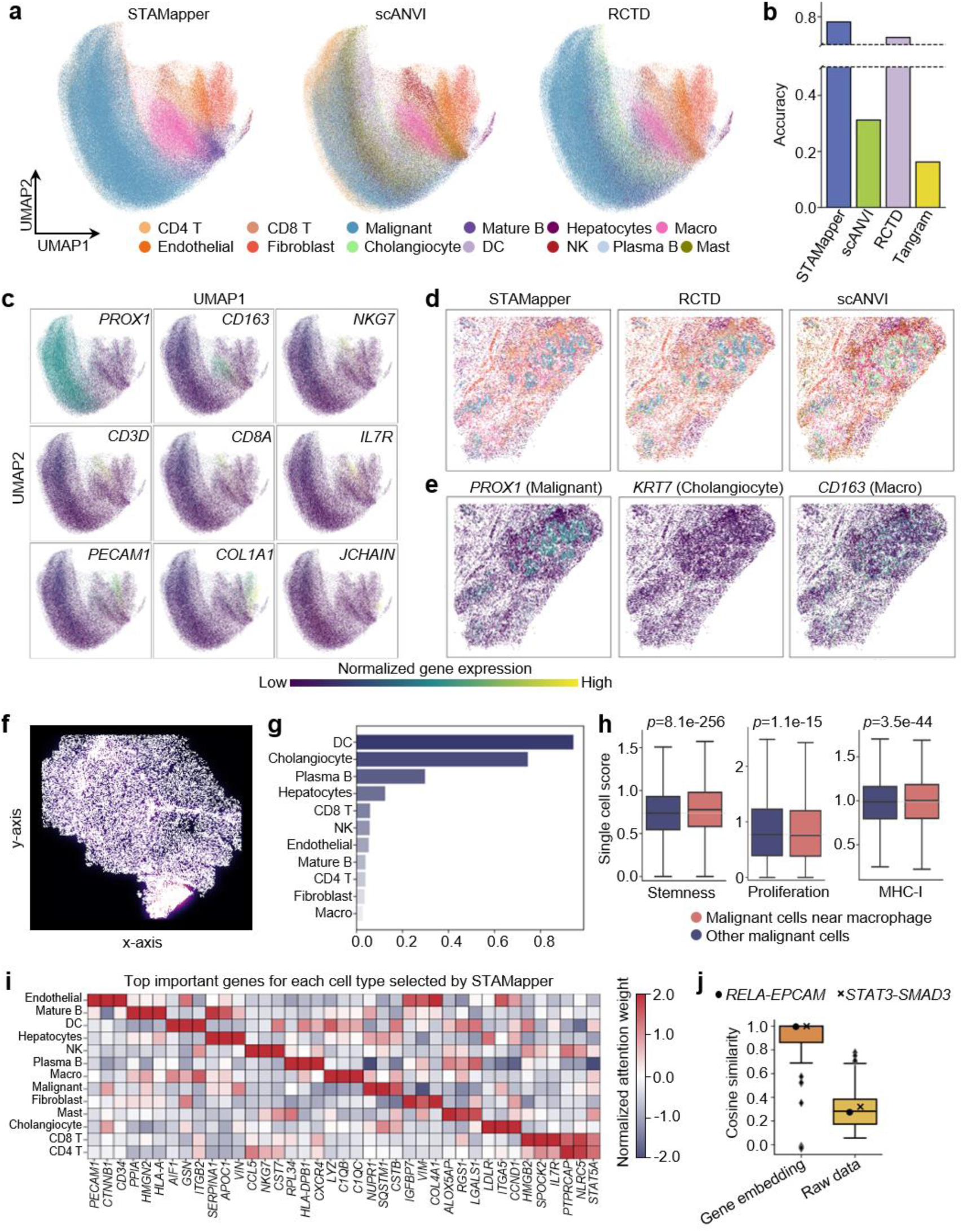
Application of STAMapper to Nanostring HCC dataset. **a.** UMAP plots of the human HCC dataset colored by STAMapper, scANVI, and RCTD, respectively. Macro, Macrophage; NK, nature killer; DC, dendritic cell. **b.** Accuracy of STAMapper, scANVI, RCTD, and Tangram on human HCC dataset. **c.** UMAP plot of the normalized marker expression corresponding to major cell types. **d.** Spatial organization of ROI 1, cells are colored by the annotation of STAMapper, RCTD, and scANVI, respectively. **e.** The normalized marker expression on ROI 1. **f.** Density plot for the distribution of macro cells. **g.** Physical distance of immune cells to malignant cells with cells being annotated by STAMapper. **h.** Boxplot for the scores of selected pathways (stemness, proliferation, and MHC-I) on malignant cells near macro and other malignant cells. **i.** Heatmap displaying genes with the highest normalized attention weights, categorized by each cell type. **j.** Cosine similarity of the gene embedding pairs learned by STAMapper, where gene pairs were TFs collected from hTFtarget. *RELA*-*EPCAM* and *STAT3-SMAD3* were validated to exist in HCC malignant cells in literature.

We found that macrophages were widely present in the sections, especially at the edges of the malignant cells, and they were the immune cells closest to the malignant cells (**Fig. 5f, g**). Macrophages are considered to play a crucial role in tumor immune evasion and also serve as pro-inflammatory mediators^32^. To investigate the role of this unique malignant-macrophage microenvironment, we classified the malignant cells into two groups, i.e., malignant cells near macrophages and others. The malignant cells near macrophages showed higher stemness^33^ and MHC-I score^34^ but lower proliferation^30^, indicating that, for malignant cells near macrophages, the increase in stemness was not due to proliferation, or they were less likely to be recognized by T cells. Therefore, it should be due to the influence of macrophages (**Fig. 5h**).

STAMapper could identify the most critical genes for each cell type and most of them are marker genes or differentially expressed genes, e.g., *PECAM1* for Endothelial^35^, *HLA-A* for Mature B^36^, *NKG7* for NK^37^, *C1QB* for Macro^38^, etc., which enhanced the model’s interpretability (**Fig. 5i**, **Methods**). Additionally, STAMapper provided gene embeddings that enhanced the similarity of functionally related gene pairs collected from hTFtarget^39, 40^. We found that two TF-target pairs, validated in the literature^41^, exhibited cosine similarities close to 1 (**Fig. 5j, Methods**). Collectively, STAMapper could provide precise annotations with interpretability, helping to discover and explain the complex tumor microenvironment.

### STAMapper reveals the cellular subpopulation localization within the layered structure of the human cortex

We applied STAMapper to the human prefrontal cortex (PFC) sequenced by Slide-tags, a whole-transcriptome single-nucleus spatial technology^11^, and the single-nucleus RNA dataset profiled for PFC samples as a reference^42^. STAMapper successfully aligned identical cell types across two datasets, including glial cells and neurons (**Fig. 6a, b**), and provided precise annotations for the scST data (**Fig. 6c**). The vascular cells (VCs, from the scRNA-seq data) and endothelial cells (ECs, from the scST data) were aligned well. That is reasonable since EC is a subtype of VC, and they both expressed *ITIH5*, a marker of ECs^11^ (**Supplementary Fig. S5a**). Utilizing the cell embeddings provided by STAMapper, we discovered a distinctive subset of oligodendrocytes (oligo) in the single-cell data that expressed *GPR17* (**Fig. 6b, d**). This gene acts as an intrinsic timer of oligodendrocyte differentiation and myelination^43^, suggesting these cells may be in a state of differentiation or development (labeled as Oligo_GPR17 for further analysis).

**Figure 6.**
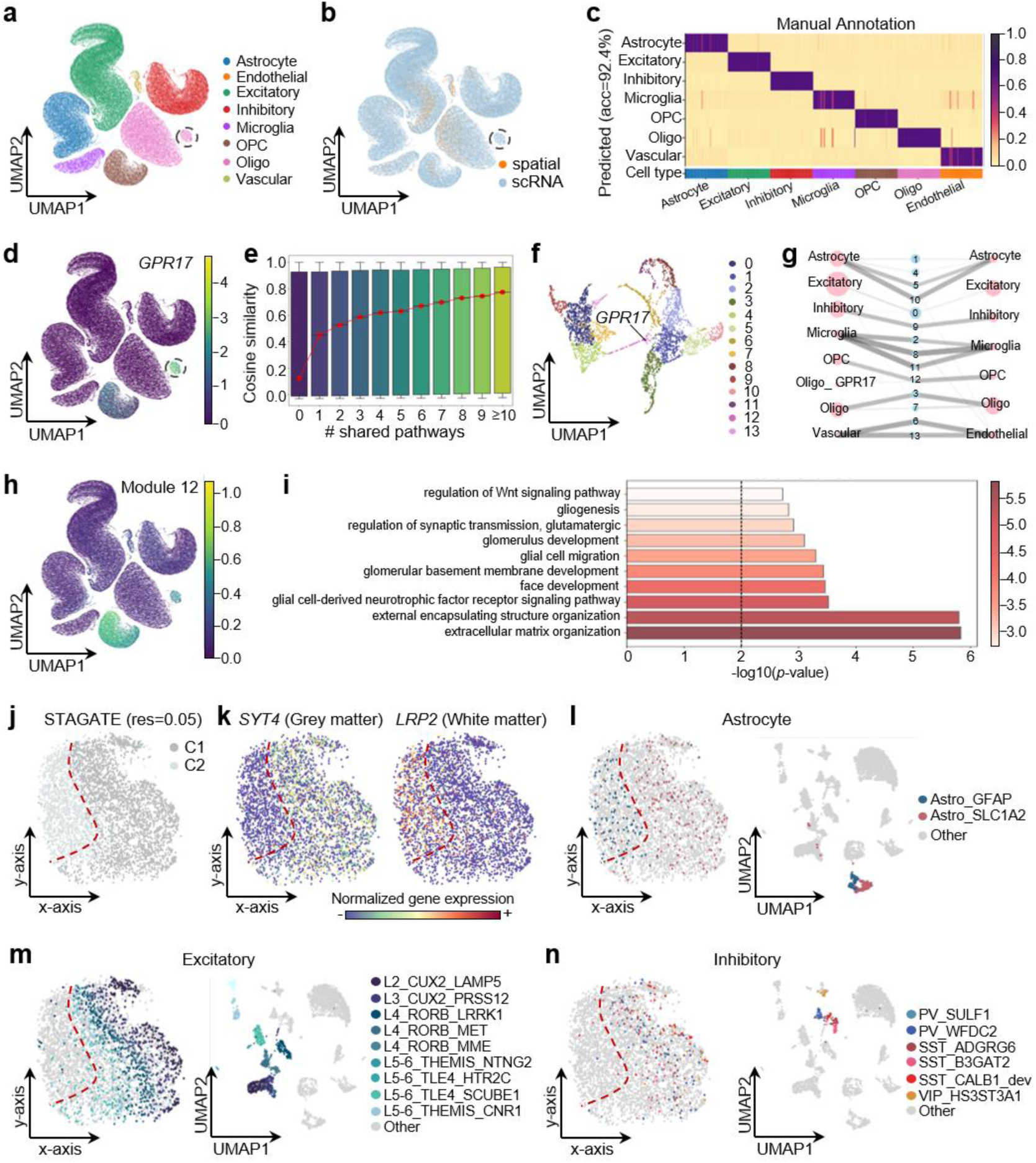
Application of STAMapper to Slide-tags human prefrontal cortex dataset. **a** and **b.** UMAP plot for the co-embedding of scRNA-seq and spatial dataset learned by STAMapper. Cells are colored based on the prediction of STAMapper (**a**) and the origin of the datasets (**b**), respectively. Oligo, oligodendrocytes; OPC, Oligodendrocyte progenitor cells. **c.** The predicted cell-type probabilities for each cell (each column) in the spatial data. A maximum of 50 cells was subsampled from each type for visualization. **d.** UMAP plots showing the co-embedding of the scRNA-seq and spatial dataset learned by STAMapper, cells are colored by the normalized expression levels of *GPR17*. **e.** Boxplots of the cosine similarity between gene embedding pairs grouped by the number of shared pathways. **f.** UMAP plot for the distribution of gene embedding, genes are colored by clusters identified through the Leiden algorithm. **g.** Abstracted graph of the heterogenous cell-gene graph, where nodes represent cell types (pink) or gene modules (blue). Node size reflects the number of cells in a cell type or genes in a module. Edge width varies with the average expression levels of cell types linked to gene modules, determined by STAMapper. **h.** UMAP plots showing the co-embedding of scRNA-seq and spatial dataset learned by STAMapper, cells are colored by the normalized expression levels of Module 12. **i**. Enrichment analysis of gene module 12 related to mouse-specific Oligo_GPR17 cells. **j.** Spatial organization of cells from the spatial dataset, cells are clustered by STAGATE with resolution=0.05. **k.** The Normalized expression of *SYT4* (marker gene of grey matter) and *LRP2* (marker gene of white matter). **l.** Spatial organization and UMAP plot of astrocyte subtypes predicted by STAMapper from the spatial dataset, subtypes with more than 20 cells are shown. The UMAP coordinates are calculated from the expression of the spatial data. **m.** Spatial organization and UMAP plot of excitatory subtypes predicted by STAMapper from the spatial dataset. Subtypes with more than 20 cells are shown. **n.** Spatial organization and UMAP plot of inhibitory subtypes predicted by STAMapper from the spatial dataset, subtypes with more than 20 cells are shown.

To further understand whether the gene embeddings provided by STAMapper captured biological meanings for each pair of input genes, we explored relationships between the number of shared pathways (Reactome^44^ and cell type signatures from MSigBD^45^) and the gene cosine similarities (**Methods**). The cosine similarities were relatively low (median=0.12) for gene pairs not presenting in any biological pathway. We observed a clear gap in the cosine similarities for gene pairs that shared a pathway (median=0.45). The more pathways shared, the higher their cosine similarities were (**Fig. 6e**). The cosine similarity achieved a median value of 0.77 for gene pairs occurring in at least ten pathways. Such a trend also existed in the raw data and was enhanced by STAMapper (**Supplementary Fig. S5b**). We next extracted 14 gene modules from these informative gene embeddings (**Fig. 6f**, **Methods**). They shared similar transcriptional patterns in the common cell types identified in both scRNA-seq and spatial data (**Fig. 6g**). Module 12 tended to be expressed in the oligodendrocyte progenitor cells (OPCs) and Oligo_GPR17 but not in oligos, and was enriched in pathways related to differentiation and development, i.e., glial cell-derived neurotrophic factor receptor signaling pathway, glial cell migration, gliogenesis and regulation of Wnt signaling pathway^46^ (**Fig. 6g-i**, **Methods**). These results further indicated that Oligo_GRP17 represents a cell type distinct from traditional oligos, potentially arising from the differentiation of OPCs.

The layered structure is a main characteristic of the cortex^47^. We next performed subtype annotations on the scST data to explore their association with the cortex structure (**Supplementary Fig. S5c, f**). To reveal the hierarchical structure of the cortex, we initially employed the clustering method STAGATE^48^ at a low resolution. STAGATE divided the section into white matter (WM) and gray matter (GM), verified by corresponding markers^49^ (**Fig. 6j, k**, **Methods**). The astrocytes expressing *GFAP* were associated with WM in mice^50^ and *SLC1A2* tended to express in the GM area^51^. STAMapper correctly annotated Astro_GFAP and Astro_SLC1A2 to the corresponding regions (**Fig. 6l**). The Oligos prefer to locate at WM^52^ (**Supplementary Fig. S5d**). On the contrary, the excitatory neurons and inhibitory located at GM are consistent with a recent study^53^ (**Fig. 6m, n**). Additionally, the subtypes of excitatory neurons exhibited layer-specific localization from the GM border to the GM/WM junction (from L2 to L6), consistent with the result of STAGATE (resolution=0.3) (**Fig. 6m**, **Supplementary Fig. S5e**). These results illustrated that STAMapper provided gene embeddings with biological meanings, facilitating the identification of shared or unique gene modules across datasets. Also, STAMapper accurately annotated structurally related cell subtypes, aiding in understanding their positional context within tissues.

## Discussion

Precise annotation of single cells from the scST data is essential for understanding the complex interactions between cellular functions and their physical locations within tissues and organs. Here, we developed an accurate and user-friendly cell-type annotation method STAMapper, which can be seamlessly integrated with the standard workflow of the package SCANPY^54^.

The primary reason for the success of STAMapper lies in its utilization of heterogeneous graph networks to simultaneously model cells and genes as two types of nodes, respectively. It employs the expression similarities among cells and the expression relationships between cells and genes. Also, it adopts a graph attention classifier to ensure that each cell pays more attention to genes that are more biologically related during the classification. However, STAMapper doesn’t incorporate the spatial information of the cells in the current version. Although it is straightforward to model the spatial information within STAMapper by connecting new edges for spatially adjacent cells, we found that this led to an accuracy loss of about 0.4% with statistical significance (**Supplementary Fig. S6a**). This could be because many spatially adjacent cells are not of the same cell type. Therefore, how to reasonably utilize spatial information to enhance the accuracy of annotations remains a challenging problem.

In this study, we concentrated on cell-type annotation for scST datasets. Given a well-annotated scRNA-seq dataset as a reference, STAMapper can accurately annotate cell types in scST data and is robust to different spatial sequencing technologies and diverse tissues. Furthermore, tests revealed that STAMapper is robust to parameter changes (**Supplementary Fig. S6b**, **Methods**). One potential limitation of STAMapper is that it utilizes only one scRNA-seq dataset as a reference, which could lead to the omission of certain cell types if the sequenced cells in the reference are not comprehensive. Future work may consider using multiple references to annotate spatial data to improve the annotation.

Selecting spatially variable genes has become a popular topic in recent years. However, the existing methods, e.g., SPARK-X^55^, STAMarker^56^, and spatialDE^57^, are designed for spatial transcriptomics technologies with lower resolution, i.e.,10X spatial transcriptomics^58^. In scST data, cell type can be employed to identify genes exhibiting spatial variation within the same cell population. STAMapper could help with an attention score to reflect the importance of each gene to a cell. If such a score undergoes a drastic change along spatial positions within a specific cell type, it could be potentially identified as a spatially variable gene relating to this cell type. We expect STAMapper to be extended in this direction.

## Methods

### Data description

We collected 81 scST datasets sequenced by different technologies, including MERFSIH, seqFISH, seqFISH+, osmFISH, STARmap, STARmap PLUS, NanoString, and Slide-tags (**Fig. 1**, **Table S1**). For the NanoString HCC data, the authors provided the annotation by an unpublished method InSituType, and we used this annotation as ground truth. For the other 80 scST datasets and all the scRNA-seq datasets, the authors provided manual annotations that served as ground truth. We also collected 16 scRNA-seq datasets sequenced by different technologies, including 10X Chromium, Droplet-microfluidic and STRT/C1 (**Fig. 1**, **Table S1**). The cell types across scRNA-seq and scST datasets were unified manually through corresponding literature and cell ontology^59^.

### Data preprocessing

In all datasets, we first normalized the library size for each cell and then logarithmized the expression data with a pseudo-count. For scRNA-seq datasets, we then selected the top 2000 highly variable genes (HVGs) and calculated the top 50 differential expressed genes (DEGs) for each cell type as input. For the scST datasets with more than 2000 genes, we used the same strategy for selecting HVGs. For the selection of DEGs, we first pre-clustered the spatial data by the Leiden algorithm^21^ (resolution=0.4) and then calculated the top 50 DEGs for each cell cluster. For the scST datasets with fewer than 2000 genes, we used the expression of all the genes as input. Finally, we scaled each gene to unit variance and zero mean value. All the preprocessing steps were implemented using the built-in functions in package SCANPY^54^.

### Construction of the heterogeneous graph

STAMapper uses a heterogeneous graph to model the two datasets with genes and cells as two types of nodes (**Fig. 1a**). We used the intersection of selected genes (DEGs and HVGs) between scRNA-seq and scST datasets as the gene nodes. We have five types of heterogeneous edges. Specifically, for each cell node and gene node, we have the edge connected to itself named ‘cell_self_loop’ and ‘gene_self_loop’. They help utilize information from the previous step in the training process. We also have ‘cell_similar_to_cell’ edges connected to similarly expressed cells with the *k* nearest neighbor strategy (based on their expression vector, *k*=5 by default) within each dataset. For a scRNA-seq dataset, we filter the edge connecting two different types of cells. Additionally, we have heterogeneous edges named ‘gene_expressed_by_cell’ and ‘cell_express_gene’ in opposite directions to indicate a gene is expressed by a cell and a cell expresses a gene.

### Architecture of STAMapper

STAMapper consists of two parts, i.e., a heterogeneous graph encoder and a heterogeneous graph attention classifier (**Fig. 1a**). STAMapper updates parameters on heterogeneous graphs according to the message-passing mechanism, where the same edge type shares the same parameters^19^. We aim to learn these edge parameters.

#### Encoder

The encoder is based on the architecture of the heterogeneous graph, where we take the expression of DEGs as the input for each cell node. Suppose we have *n* cells in the scRNA-seq dataset and *m* cells in the scST dataset.

Here, we denote 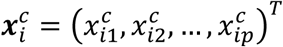 as the input for cell node *i* (*i* ∈ {1, …, *n*, …, *N* = *m* + *n*}, where 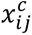 denotes the normalized expression for gene *j* (*j* ∈ {1,2, …, *p*}) in cell *i*. For the cell node *i* the initial embedding is calculated as follows:

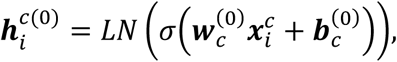

where *LN* denotes the layer normalization^60^, *σ* is the nonlinear activation function, 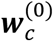 denotes the learnable parameters for the edge type ‘cell_self_loop’ in the initial layer, 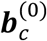 denotes the learnable bias for cell nodes. We used the intersection of selected genes (DEGs and HVGs) between scRNA-seq and scST datasets as the gene nodes. Suppose we have *s* gene nodes, the initial embedding for gene node *k* is as follows:

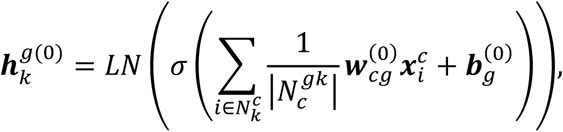

where 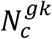 is a set containing the cell nodes which are neighbors of gene node *k* where 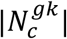 denotes the number of cells in this set. 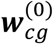 denotes the learnable parameters for edge type ‘cell_express_gene’ for the initial layer, 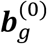 denotes the learnable bias for gene nodes.

For the *l*-th (1 ≤ *l* ≤ *L*) hidden layer, the embedding for cell node *i* is:

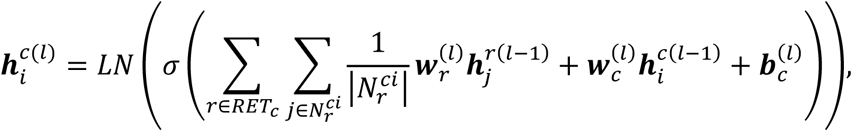

where *RET*_*c*_ denotes a set of edge types connected to node *c*, 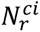 denotes a set containing the nodes that are neighbors of cell node *i* according to edge type *r*, 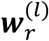 denotes the learnable parameters for edge type *r*, 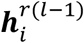 denotes the embedding for the *i*-th cell/gene node (adaptive to the edge type *r*) in the previous layer. The embedding for gene node *k* in the *l*-th hidden layer is:

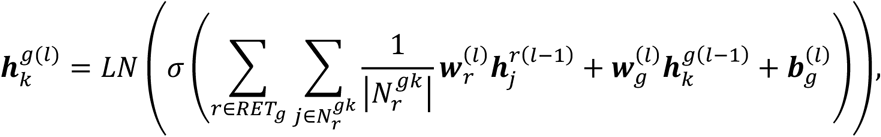

*RET*_*g*_ denotes a set of edge types connecting to node *g*, 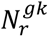 denotes a set containing the nodes that are neighbors of gene node *k* according to edge type *r*, 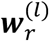 denotes the learnable parameters for edge type *r*, 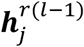 denotes the embedding for the *j*-th cell/gene node (adaptive to the edge type *r*) in the previous layer.

#### Classifier

To further utilize the information from genes associated with cell classification, we employed an attention mechanism^19^ in the heterogeneous graph classifier. Specifically, the attention weight of the classifier from gene node *j* to cell node *i* is:

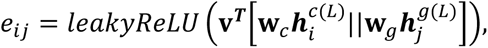

Where **v** is a learnable weight vector, 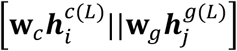 denotes concatenation of 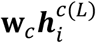 and 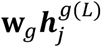. The weight is further normalized as follows:

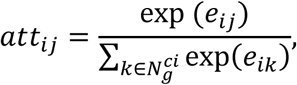

where 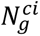 is a set containing the gene nodes that are neighbors of cell node *i*. Then the output logits of the classifier are:

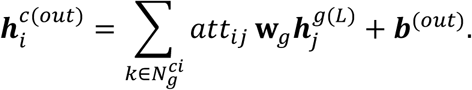

Then we apply *softmax* function over logits coordinately:

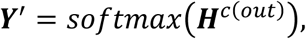

where 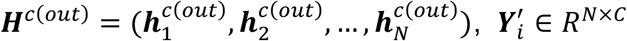. Here, *N* denotes the total number of cells, ***C*** denotes the number of cell types, 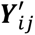 denotes the predicted probabilities of cell type *j* for the *i*-th cell.

#### Loss function

We modified the cross-entropy loss as the classification loss. Suppose ***y***_*sc*_ is the manual annotation of cells from the scRNA-seq dataset. We first use a weighted cross-entropy loss as follows:

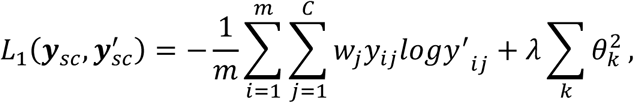

Where 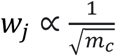 denotes the weight for cell-type *j*, *m*_*c*_ denotes the number of cells in *c*-th cell type, 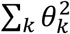 denotes the sum of the squares of all model parameters penalizing the complexity of the model to prevent overfitting, and *λ* is set as 0.01 by default. We applied the label smoothing technique to reduce the model overconfidence^20^. The overall training loss is:

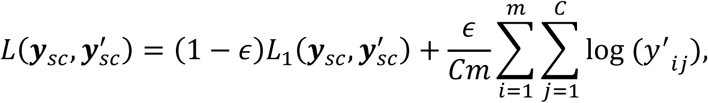

the default value of *ϵ* is 0.1.

#### Training process

In all experiments, we set the encoder of STAMapper as a two-layer heterogeneous graph neural network with 512 hidden units. We apply the Adam optimizer^61^ with a learning rate of 5e-3 to optimize all the parameters. We adopt LeakyReLU^62^ as the activation function with a negative slope set as 0.05. We set the number of iterations as 1000 by default and use the same checkpoint selection strategy as a recently published method^63^.

### Unknown cells detection

We detected unknown cells based on two criteria: (i) cells in the scST data with predicted probability less than the median value; and (ii) cell types with distance from the scST data to scRNA-seq data larger than a user-defined threshold (default 4). The distance was calculated based on the cell embeddings provided by STAMapper, after reducing to 50 dimensions using PCA. Specifically, the distance for a given cell type from the scST data to scRNA-seq data is defined as the average distance of each cell in the scST data to its five nearest neighbors in the scRNA-seq data.

### Gene module extraction and enrichment analysis

We performed the Leiden community detection algorithm^21^ on the gene embeddings from the last layer of STAMapper and defined the detected clusters as gene modules. We performed the GO enrichment analysis by the R package clusterProfiler^64^ with the biological process ontologies.

### Benchmarking cell-type annotation

We performed scANVI^15^ with the Python package scVI, referred to the section “Integration and label transfer with Tabula Muris” from its documentation (https://docs.scvi-tools.org/en/stable/tutorials/notebooks/scrna/tabula_muris.html). We performed RCTD using the R package spacexr^16^ with default workflow (https://raw.githack.com/dmcable/spacexr/master/vignettes/spatial-transcriptomics.html). We performed Tangram^17^ with the Python package with cluster mode (https://tangram-sc.readthedocs.io/en/latest/tutorial_sq_link.html).

### Spatial clustering with STAGATE

We performed STAGATE^48^ on the Slide-tags cortex data and followed a standard workflow for data preprocessing with a rad_cutoff as 200 to construct the spatial network (https://stagate.readthedocs.io/en/latest/T3_Slide-seqV2.html).

### Data availability

All datasets analyzed in this study are publicly available. The raw datasets are available from the following studies:

**Dataset 1-4 (mouse prefrontal cortex):** STARmap, https://github.com/weallen/STARmap/tree/master; 10X Chromium, adult samples from GSE124952 in the GEO database.

**Dataset 5-8 (mouse visual vortex):** STARmap, https://github.com/weallen/STARmap/tree/master; Smart-seq, https://portal.brain-map.org/atlases-and-data/rnaseq/mouse-v1-and-alm-smart-seq.

**Dataset 9 (mouse visual vortex):** seqFISH+, https://github.com/CaiGroup/seqFISH-PLUS; Smart-seq, https://portal.brain-map.org/atlases-and-data/rnaseq/mouse-v1-and-alm-smart-seq.

**Dataset 10 (mouse somatosensory cortex):** osmFISH, https://github.com/drieslab/spatial-datasets/tree/master/data/2018_osmFISH_SScortex/raw_data; STRT/C1, GSE60361 in the GEO database.

**Dataset 11-23 (mouse primary motor cortex):** MERFISH, https://knowledge.brain-map.org/data/L3GYGFMDJCG0GUEE3QG/collections; 10X Chromium, https://data.nemoarchive.org/biccn/lab/zeng/transcriptome/scell/10x_v3/mouse/processed/analysis/10X_cells_v3_AIBS/. **Dataset 24-33 (mouse retina):** MERFISH, https://doi.org/10.5281/zenodo.8144355; 10X Chromium, GSE135406 in the GEO database.

Dataset 34 (mouse kidney): MERFISH, https://figshare.com/projects/MERFISH_mouse_comparison_study/134213; 10X Chromium, https://figshare.com/articles/dataset/SingleCellData_raw_/19310675.

**Dataset 35 (mouse liver):** MERFISH, https://figshare.com/projects/MERFISH_mouse_comparison_study/134213; 10X Chromium, https://figshare.com/articles/dataset/SingleCellData_raw_/19310675.

**Dataset 36-71 (mouse hypothalamic preoptic region):** MERFISH, https://datadryad.org/stash/dataset/doi:10.5061/dryad.8t8s248/; Droplet-microfluidic, GSE113576 in the GEO database.

**Dataset 72-77 (mouse gastrulation):** osmFISH, seqFISH, https://marionilab.cruk.cam.ac.uk/MouseGastrulation2018/; 10X Chromium, https://bioconductor.org/packages/devel/data/experiment/vignettes/MouseGastrulationData/inst/doc/MouseGastrulationData.html.

**Dataset 78-79 (mouse prefrontal cortex):** STARmap PLUS, https://github.com/weallen/STARmap/tree/master; 10X Chromium, GSE124952 in the GEO database.

**Dataset 80 (human prefrontal cortex):** Slide-tags, https://singlecell.broadinstitute.org/single_cell/study/SCP2167/slide-tags-snrna-seq-on-human-prefrontal-cortex#study-download; 10X Chromium, GSE168408 in the GEO database.

**Dataset 81 (human liver cancer):** NanoString, https://nanostring.com/resources/liver-cancer-raw-data-files-cosmx-smi-human-liver-ffpe-dataset/; 10X Chromium, GSE149614 in the GEO database.

### Code availability

STAMapper is implemented in Python and is available on GitHub [https://github.com/zhanglabtools/STAMapper].

## Acknowledgments

This work has been supported by the National Key Research and Development Program of China (nos. 2019YFA0709501 to S.Z. and 2021YFC270160 to S.Z.), the Science and Technology Commission of Shanghai Municipality (no. 23JC1401000 to S.Z.), the National Natural Science Foundation of China (nos. 32341013, 12326614, 12126605 to S.Z.), the R&D project of Pazhou Lab (Huangpu) (no. 2023K0602 to S.Z.) and the CAS Project for Young Scientists in Basic Research (no. YSBR-034 to S.Z.).

## Author contributions

Shihua Zhang conceived and supervised the project. Qunlun Shen designed and implemented the STAMapper algorithm. Qunlun Shen and Kangning Dong validated the study. Qunlun Shen, Shuqin Zhang and Shihua Zhang wrote the manuscript. All authors read and approved the final manuscript.

## Supplementary Materials

### Supplementary Figures

**Supplementary Figure S1.**
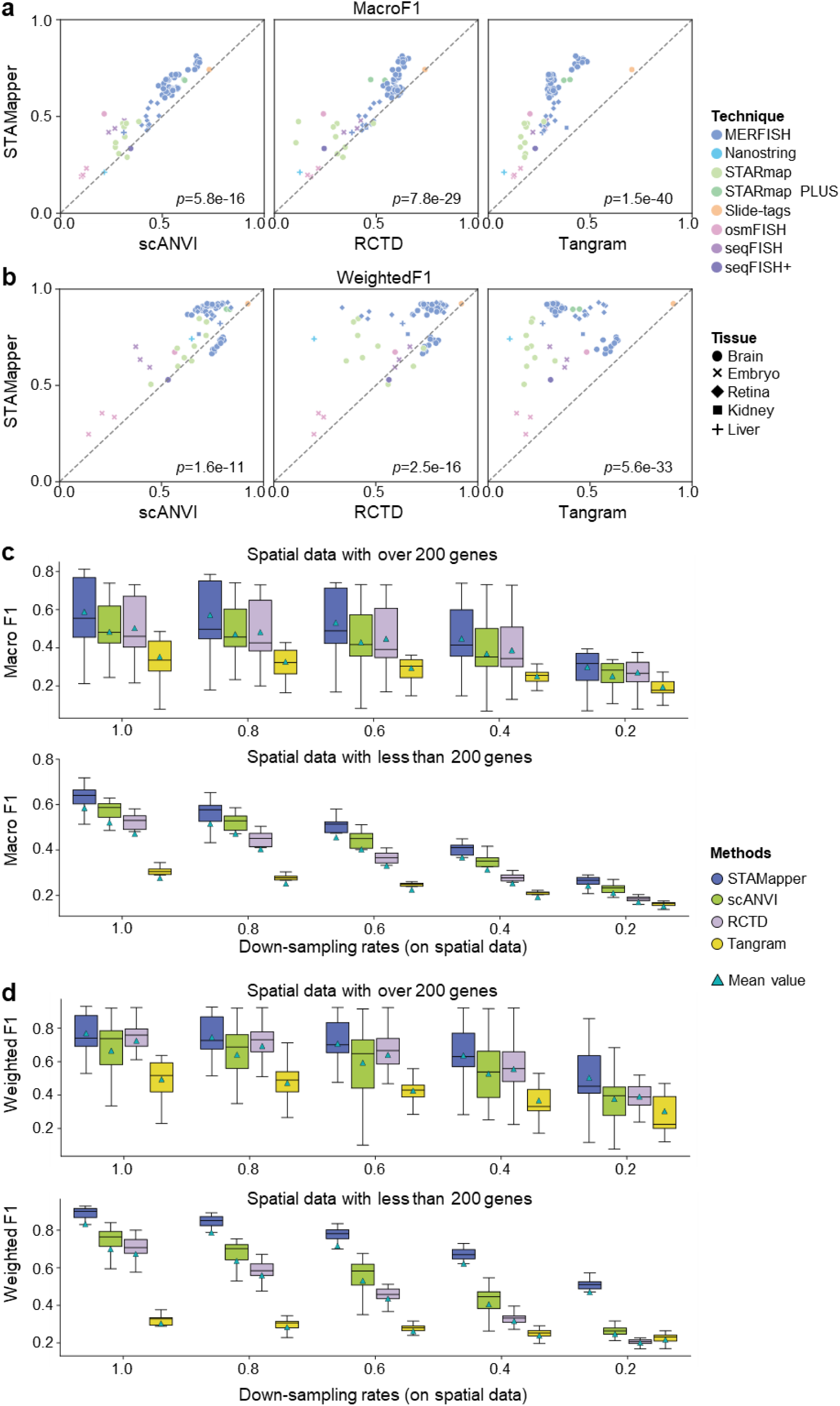
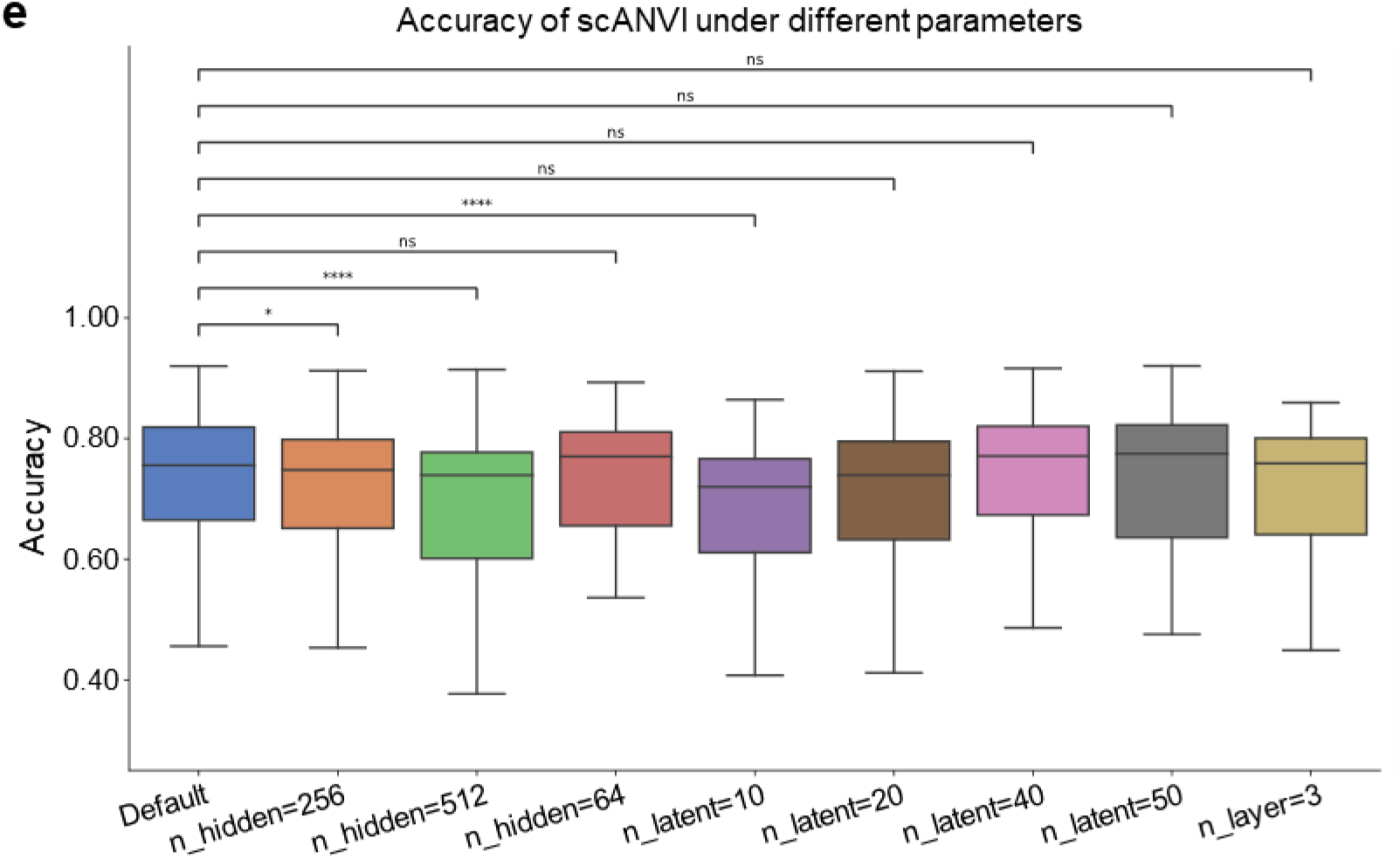
**a** and **b.** Performance comparison of STAMapper and three other methods in terms of macro F1 score **(a)** and weighted F1 score **(b)** on the 81 pairs of scRNA-seq and scST datasets. *p* values were calculated by paired t-test. **c** and **d.** Performance comparison of the macro F1 score **(c)** and weighted F1 score **(d)** of STAMapper and three other methods on different down-sampling rates (1.0, 0.8, 0.6, 0.4, 0.2) for read counts, where the down-sampling rate of 1.0 means the raw data. The upper panel depicts the scST datasets with more than 200 genes for sequencing, while the lower panel corresponds to fewer than 200 genes. **e.** The accuracy of scANVI under different parameters, with the default settings being n_hidden=128, n_layers=2, and n_latent=30.

**Supplementary Figure S2.**
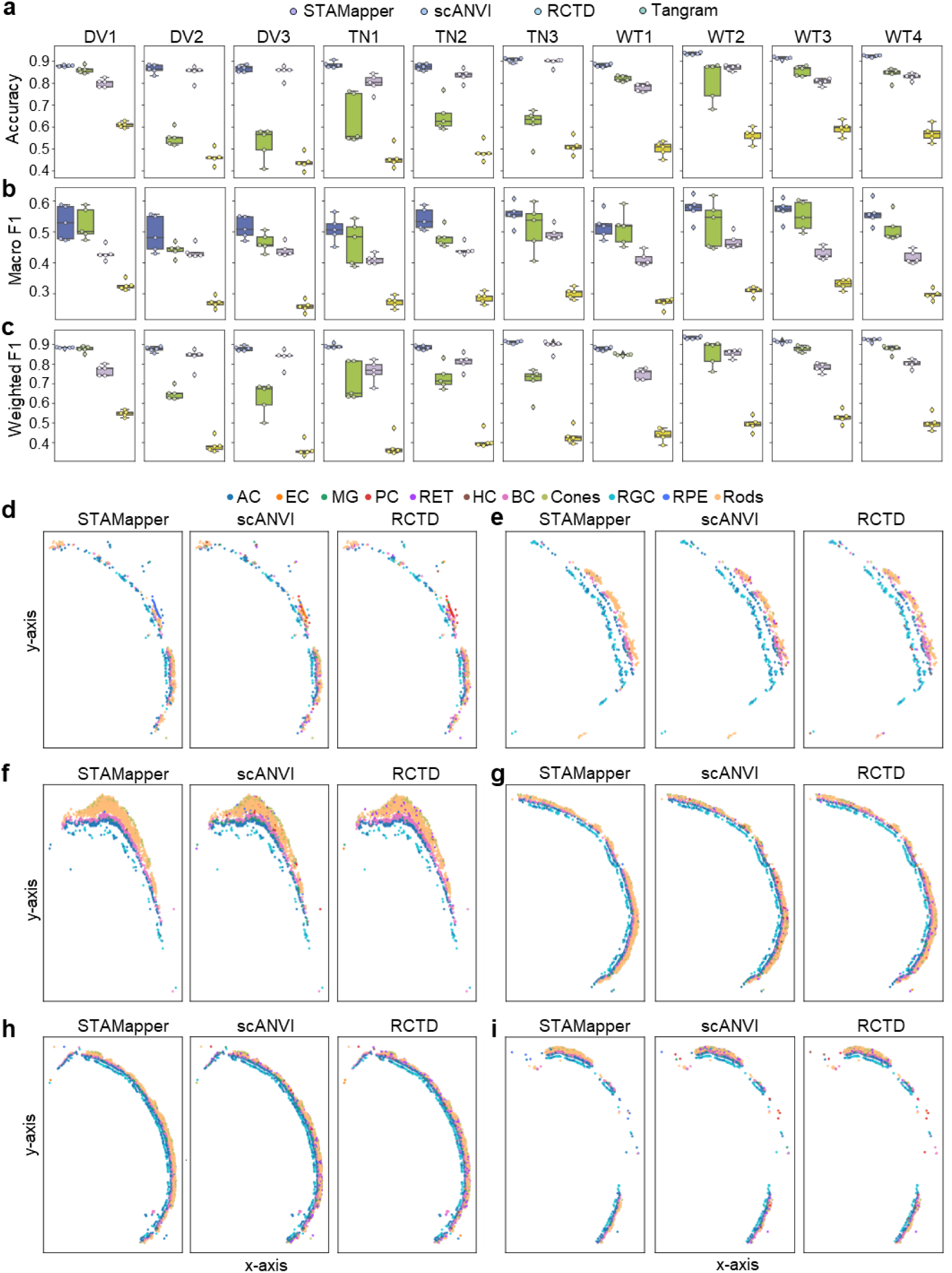
**a** and **c.** Box plots showing the accuracy **(a)**, macro F1 score **(b)**, and weighted F1 score **(c)** on the 50 paired datasets (5 scRNA-seq datasets and 10 scST datasets). Each column corresponds to a scST dataset. **d-i.** Spatial organization of the remaining six slices from the scST data corresponding to Figure 2b, where cells are colored by the annotation by STAMapper, scANVI, and RCTD, respectively.

**Supplementary Figure S3.**
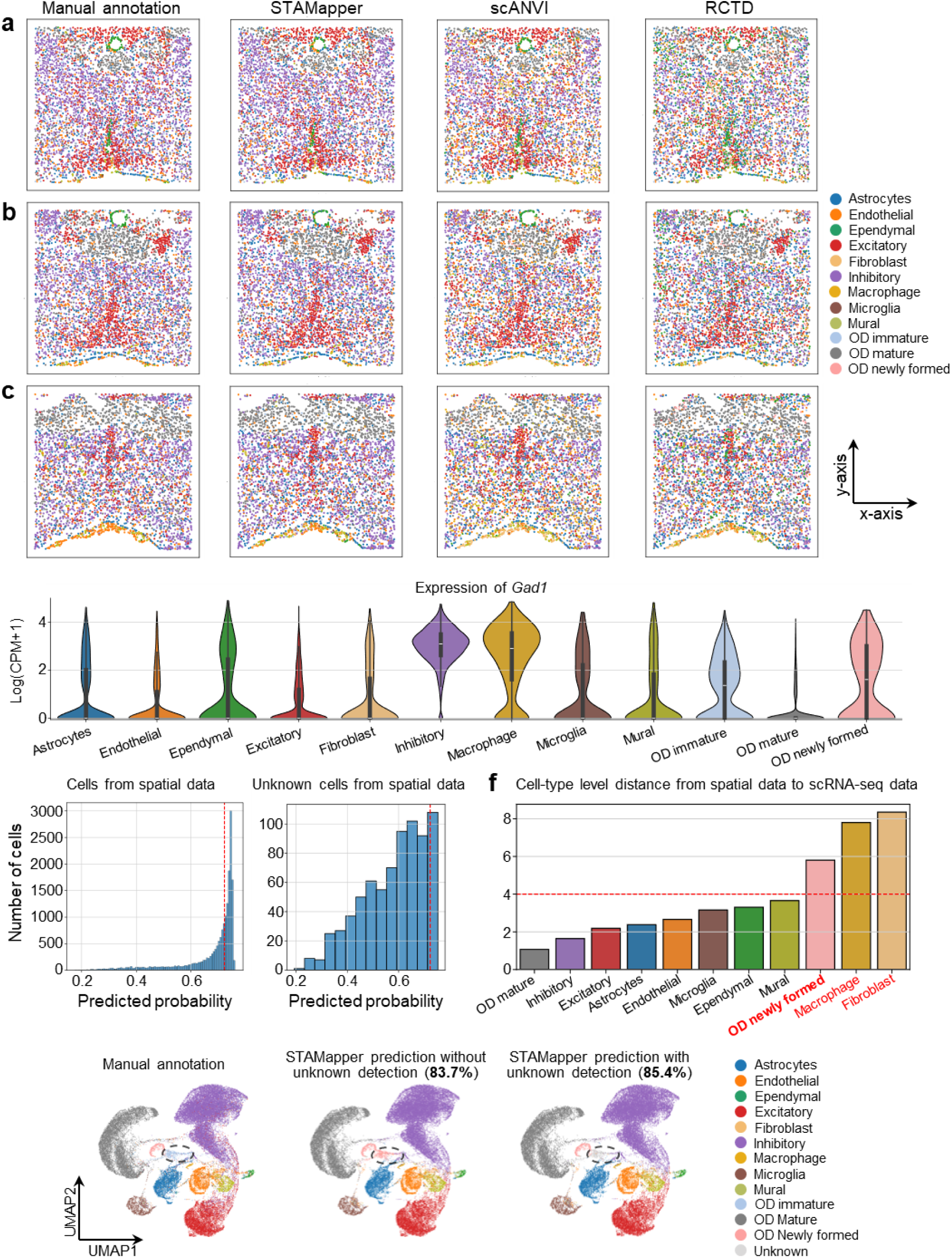
**a-c**, Spatial organization of the remaining three slices from the mouse hypothalamic data corresponding to Figure 4b. The cells are colored based on the manual annotation and the prediction by STAMapper, scANVI, and RCTD, respectively. **d.** Expression levels of *Gad1* (a marker of inhibitory cells) across different cell types (annotated by scANVI). **e.** The predicted probability of STAMapper for each cell from the scST data (left panel), the predicted probability of STAMapper for unknown cells from the scST data (right panel). **f.** Cell-type level distance from the scST data to the scRNA-seq data on the embedding learned by STAMapper. Bold font indicates unknown cells were predicted as this specific cell type, and red font denotes cell types present in single-cell data but not annotated in the scST data by manual annotation. **g.** UMAP plots showing the co-embedding of the scRNA-seq and scST data learned by STAMapper, where cells are colored by manual annotation, STAMapper prediction without unknown detection, and STAMapper prediction with unknown detection, respectively. The percentages in parentheses represent the predicted accuracy.

**Supplementary Figure S4.**
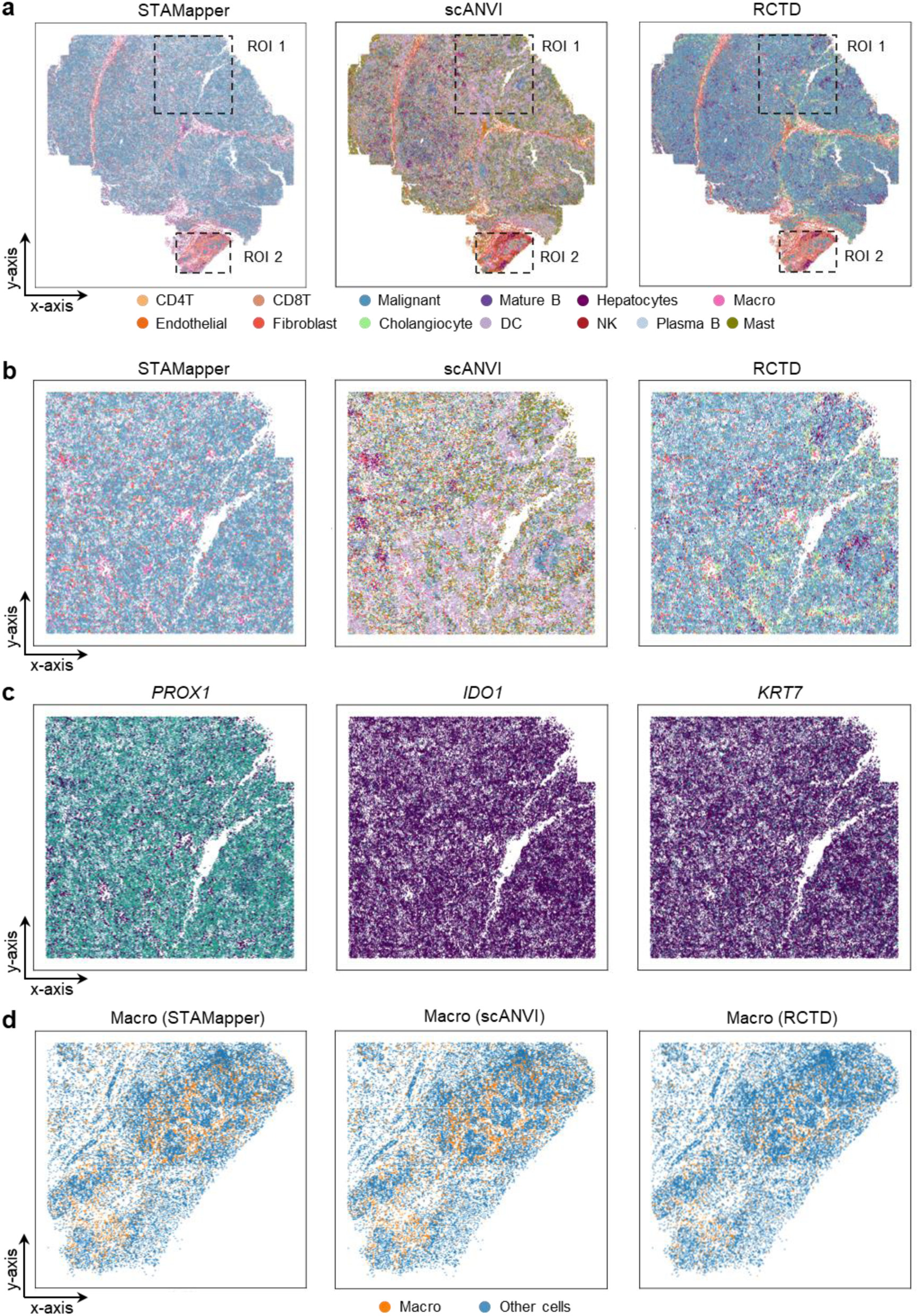
**a.** Spatial organization of the Nanostring HCC dataset corresponding to Figure 5a. **b.** Spatial organization of ROI 1, where cells are colored based on the annotation by STAMapper, RCTD, and scANVI, respectively. **c** and **d.** The normalized maker expression on ROI 1 **(c)** and ROI 2 **(d)**.

**Supplementary Figure S5.**
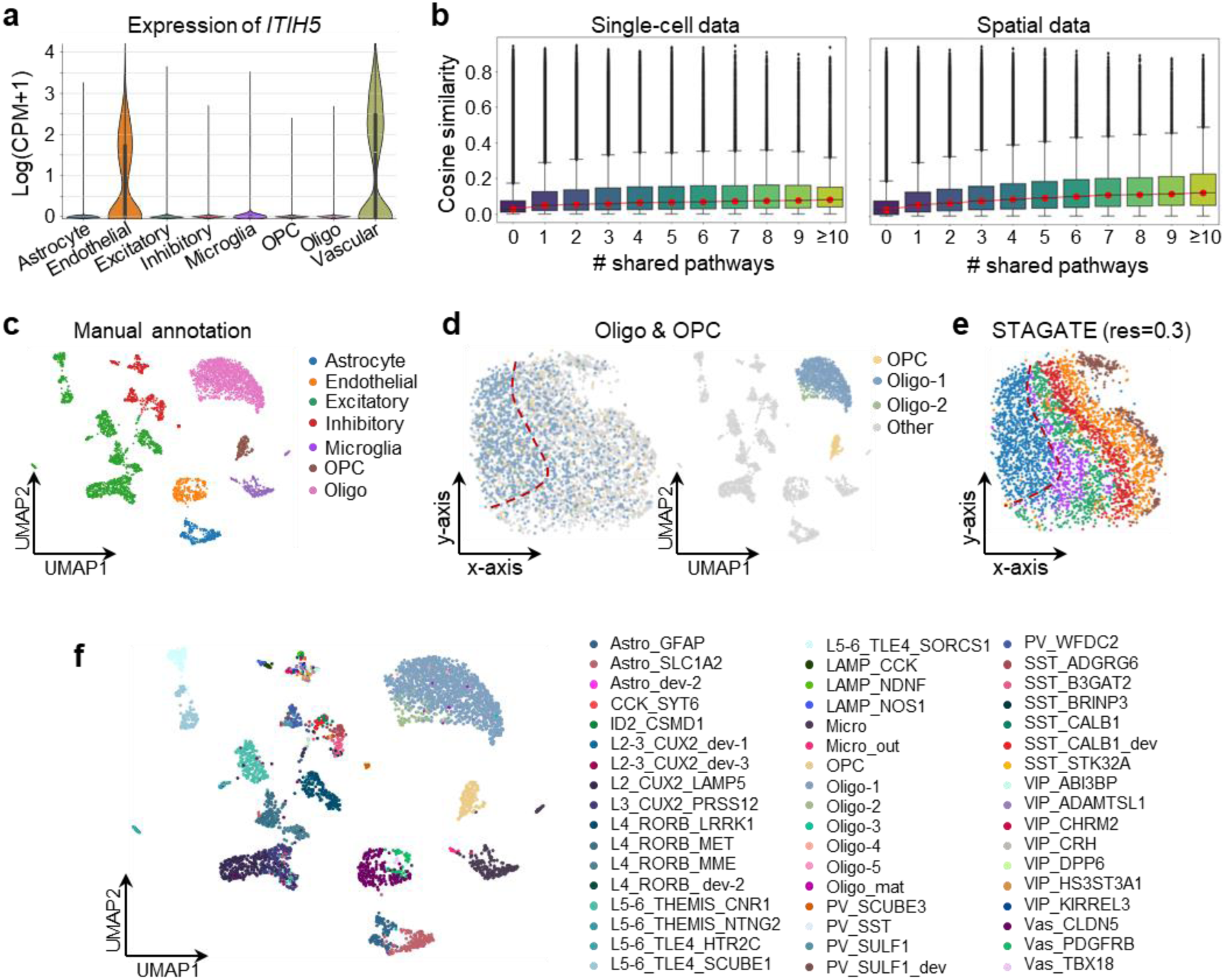
**a.** Violin plot showing the expression of *Sema4d* across different cell types. **b.** Boxplots showing the cosine similarity between the expression vector of gene pairs from the scRNA-seq data (left panel) and scST data (right panel), and the gene pairs are grouped by the number of shared pathways. **c.** UMAP plots of the spatial data and cells colored by manual annotation. **d.** Spatial organization and UMAP plot of Oligo & OPC subtypes (predicted by STAMapper) from the scST data and subtypes with more than 20 cells are shown. **e.** Spatial organization of cells from the scST data, where cells are clustered by STAGATE with resolution=0.3. **f.** UMAP plots showing the distribution of cell subtypes on the scST data learned by STAMapper.

**Supplementary Figure S6.**
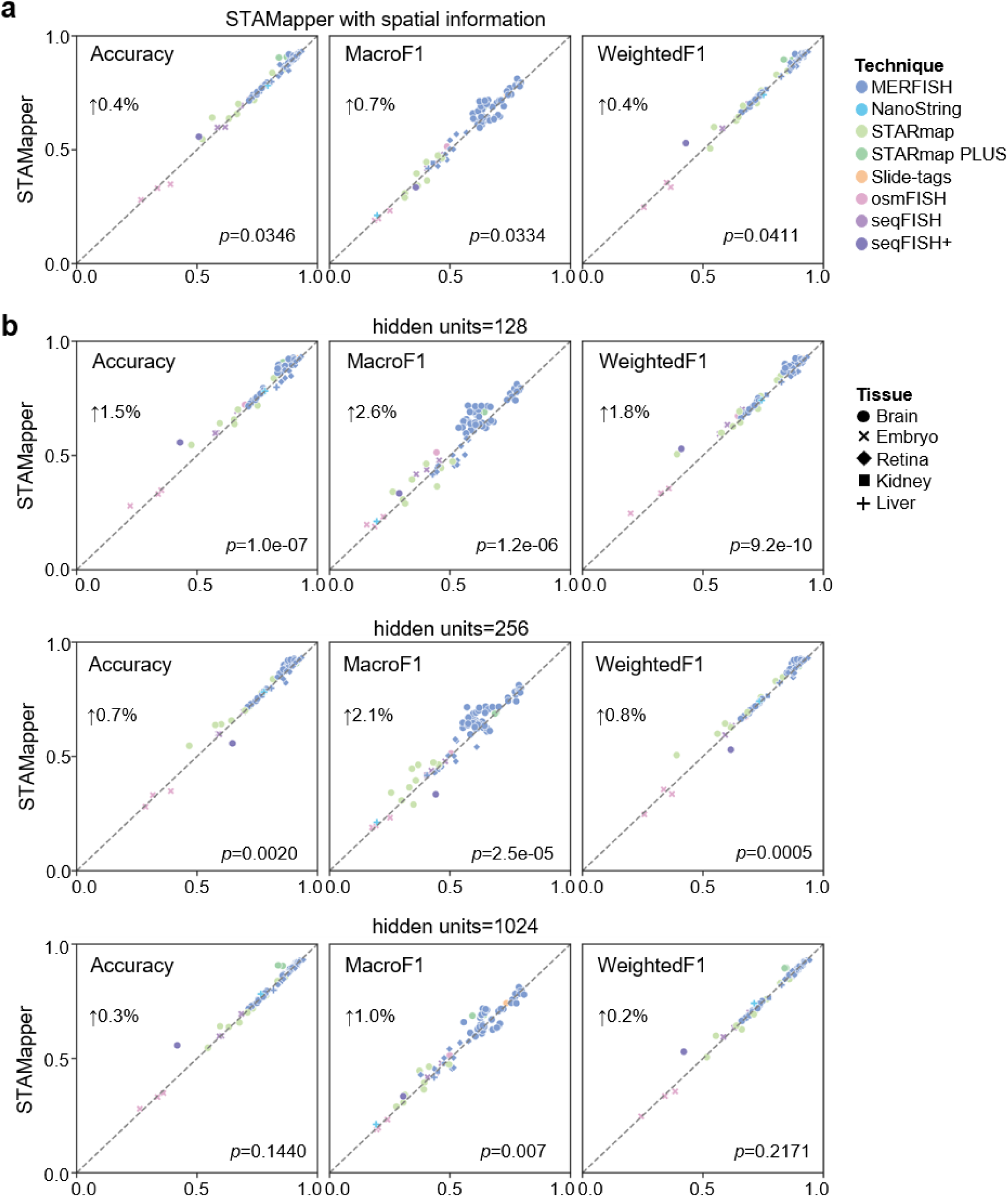
**a.** Performance comparison of STAMapper (default parameters) and STAMapper with spatial information regarding cell-annotation accuracy, macro F1 score and weighted F1 score on 81 pairs of scRNA-seq and scST datasets. *P*-values were calculated by paired t-test. **b.** Performance comparison of STAMapper (default parameters) and STAMapper under different numbers of hidden units (i.e., 128, 256 and 1024) regarding cell-annotation accuracy, macro F1 score and weighted F1 score on 81 pairs of scRNA-seq and scST datasets. *P*-values were calculated by paired t-test.

### Supplementary Tables

**Table S1.** Description of all scRNA-seq datasets used in this study.

| Technique | Tissue | Description | #Genes | #Data sets | Reference |
| --- | --- | --- | --- | --- | --- |
| **Droplet-microfluidic** | Mouse brain | Hypothalamic preoptic region | 20320 | 1 | 1 |
| **10X Chromium** | Mouse kidney | Kidney | 20138 | 1 | 2 |
| **10x Chromium** | Mouse liver | Liver | 20138 | 1 | 2 |
| **Smart-seq** | Mouse brain | Visual vortex | 34041 | 1 | 3 |
| **10X Chromium** | Mouse brain | Prefrontal cortex | 19517 | 1 | 4 |
| **10X Chromium** | Mouse brain | Primary motor cortex | 26523 | 1 | 5 |
| **STRT/C1** | Mouse brain | Somatosensory cortex | 19972 | 1 | 6 |
| **10X Chromium** | Mouse retina | Retina | 27998 | 5 | 7 |
| **10X Chromium** | Mouse embryo | Gastrulation | 19362 | 1 | 8 |
| **10X Chromium** | Mouse brain | Prefrontal cortex | 21000 | 1 | 4 |
| **10X Chromium** | Human brain | Prefrontal cortex | 26747 | 1 | 9 |
| **10X Chromium** | Human liver | Hepatocellular Carcinoma | 25479 | 1 | 10 |

**Table S2.** Description of all scST datasets used in this study.

| **Technique** | **Tissue** | **Description** | **#Genes** | **#Data sets** | **Reference** |
| --- | --- | --- | --- | --- | --- |
| **STARmap** | Mouse brain | Prefrontal cortex | 166 | 4 | 11 |
| **STARmap** | Mouse brain | Visual vortex | 166 | 3 | 11 |
| **STARmap** | Mouse brain | Visual vortex | 1020 | 2 | 11 |
| **seqFISH+** | Mouse brain | Visual vortex | 10000 | 1 | 12 |
| **osmFISH** | Mouse brain | Somatosensory cortex | 33 | 1 | 13 |
| **MERFISH** | Mouse brain | Primary motor cortex | 254 | 12 | 14 |
| **MERFISH** | Mouse retina | Retina | 368/500 | 10 | 15 |
| **MERFISH** | Mouse kidney | Kidney | 307 | 1 | 2 |
| **MERFISH** | Mouse liver | Liver | 307 | 1 | 2 |
| **MERFISH** | Mouse brain | Hypothalamic preoptic region | 161 | 36 | 1 |
| **osmFISH** | Mouse embryo | Gastrulation | 33 | 3 | 8 |
| **seqFISH** | Mouse embryo | Gastrulation | 251 | 3 | 8 |
| **STARmap PLUS** | Mouse brain | Prefrontal cortex | 2766 | 2 | 16 |
| **Slide-tags** | Human brain | Prefrontal cortex | 36601 | 1 | 17 |
| **NanoString** | Human liver | Hepatocellular Carcinoma | 1000 | 1 | 18 |

